# MultiHeadGAN: A Deep Learning Method for Low Contrast Retinal Pigment Epithelium Cells Segmentation in Fluorescent Flatmount Microscopy Images

**DOI:** 10.1101/2022.03.29.486292

**Authors:** Hanyi Yu, Fusheng Wang, George Theodoro, John Nickerson, Jun Kong

**Affiliations:** Department of Computer Science, Emory University, Atlanta, GA; Department of Computer Science, Stony Brook University, Stony Brook, NY; Department of Computer Science, Federal University of Minas Gerais, Belo Horizonte, Brazil; Department of Ophthalmology, Emory University, Atlanta, GA; Department of Mathematics and Statistics, Georgia State University, Atlanta, GA

**Keywords:** Deep learning, Retinal pigment epithelium, Image segmentation, Generative adversarial networks, Semi-supervised learning

## Abstract

**Background:** Retinal pigment epithelium (RPE) aging is an important cause of vision loss. As RPE aging is accompanied by changes in cell morphological features, an accurate segmentation of RPE cells is a prerequisite to such morphology analyses. Due the overwhelmingly large cell number, manual annotations of RPE cell borders are time-consuming. Computer based methods do not work well on cells with weak or missing borders in the impaired RPE sheet regions.

**Method:** To address such a challenge, we develop a semi-supervised deep learning approach, namely MultiHeadGAN, to segment low contrast cells from impaired regions in RPE flatmount images. The developed deep learning model has a multi-head structure that allows model training with only a small scale of human annotated data. To strengthen model learning effect, we further train our model with RPE cells without ground truth cell borders by generative adversarial networks. Additionally, we develop a new shape loss to guide the network to produce closed cell borders in the segmentation results.

**Results:** In this study, 155 annotated and 1,640 unlabeled image patches are included for model training. The testing dataset consists of 200 image patches presenting large impaired RPE regions. The average RPE segmentation performance of the developed model MultiHeadGAN is 85.4 (correct rate), 88.8 (weighted correct rate), 87.3 (precision), and 80.1 (recall), respectively. Compared with other state-of-the-art deep learning approaches, our method demonstrates superior qualitative and quantitative performance.

**Conclusions:** Suggested by our extensive experiments, our developed deep learning method can accurately segment cells from RPE flatmount microscopy images and is promising to support large scale cell morphological analyses for RPE aging investigations.

## Introduction

The retinal pigment epithelium (RPE) is a pigmented cell layer between the choroid and the neurosensory retina. Among others, the main RPE functions are to transport nutrients, maintain the photoreceptor excitability, and secrete immunosuppressive factors [1]. Aging of the RPE can cause the loss or reduction of the indicated functions that affect the function and survival of photoreceptor cells and choroidal cells. Therefore, it may result in the secondary degeneration of photoreceptors and finally lead to irreversible vision loss [2]. Previous studies have suggested that RPE cell morphological features, such as area, perimeter, aspect ratio, polymegathism, and pleomorphism, can be indicators of the cell pathophysiologic status to determine the degree of RPE aging [3–5].

RPE flatmount images have been widely used to calculate RPE cell morphological features. An accurate detection of cell borders from flatmount microscopy images is a prerequisite for determining cell morphological features. In our prior work [5], a machine-learning-based ImageJ (National Institutes of Health, Bethesda, MD, USA) plugin known as Trainable Weka Segmentation [6] was utilized to extract cell borders with a limited success, especially in impaired regions enriched with weak or missing RPE cell borders. Typical examples of damaged regions are given in Figure 1. To ensure the accuracy of downstream morphology analyses, manual post processing steps are, therefore, required to remove these damaged regions from further analyses. This process is not only time-consuming, but also significantly reduces the scale of data for analysis, resulting in a weaker study power. More importantly, it makes infeasible to study RPE cell morphology and structures within impaired regions necessary for RPE recovery mechanism and aging investigations. Thus, it is imperative to develop an effective and efficient approach to recover blurred and missing RPE cell borders in large scale flatmount microscopy images.

**Figure 1.**
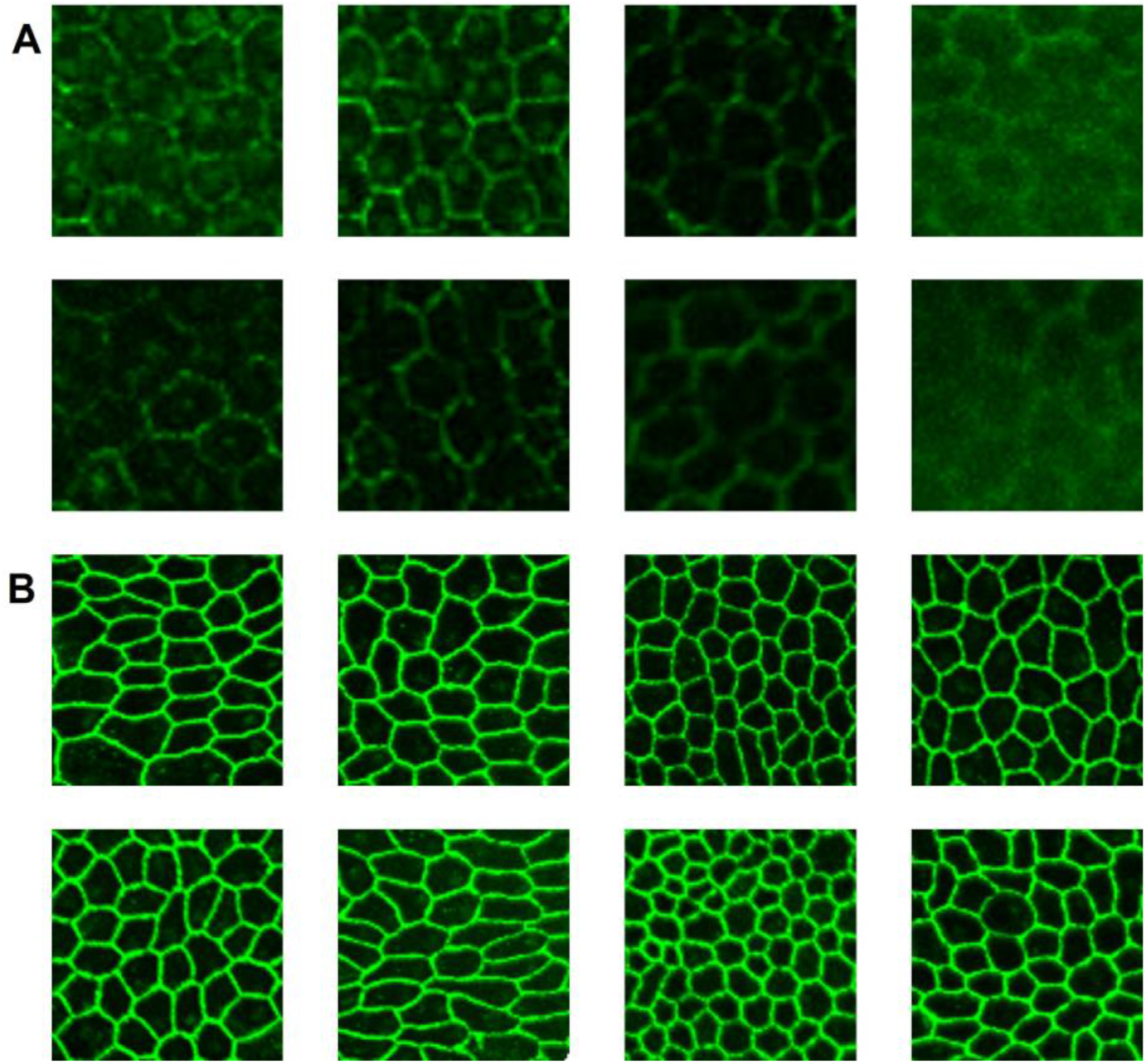
Representative examples of RPE flatmount image regions. (A) RPE cells in damaged regions present weak or missing cell borders with partial or complete cell structure loss. (B) RPE cells in normal regions often have cells with high contrast borders.

Unlike traditional machine learning methods, deep neural networks require no manual feature engineering and present an enhanced data learning power to support biomedical research [7,8]. The basic structure of a deep neural network is composed of layers of computational nodes analogous to brain neurons. For semantic segmentation tasks, a class of deep neural networks consists of an encoder for latent feature extraction from input images and a decoder for mapping the extracted features to desired segmentation results. Deep neural networks have been widely used for segmentation with multiple image modalities, ranging from bright field histopathology image slides [9,10], CT [11,12], MRI [13], and immunofluorescence microscopy images [14,15]. Although there are multiple state-of-the-art deep learning models [15–18] that can be potentially used to segment RPE cells presenting blurred or missing cell borders, they require a large scaled annotated training dataset. As the manual annotations on RPE cells in damaged regions are time-consuming, we only have a small set of annotated weak RPE cells insufficient to support the supervised learning strategy by these state-of-the-art deep learning models. To address this challenge, we leverage the Generative Adversarial Networks (GAN) mechanism to enrich the training dataset with a massive amount of unlabeled weak RPE cells and mitigate the model overfitting problem. The resulting deep learning model, namely MultiHeadGAN, is built upon the state-of-the-art UNet, but with a new training strategy simultaneously leveraging a small set of annotated and a large set of unlabeled RPE cells from flatmount microscopy images for morphology feature extraction and RPE structure reconstruction. Additionally, we design a new shape loss for model training that favors closed cell borders. We present the efficacy of our model design by ablation experiments. Our method is both qualitatively and quantitatively evaluated and compared with the state-of-the-art deep learning approaches. The extensive experimental results demonstrate the superiority of our developed segmentation method, suggesting its potential to facilitate further biomedical research on RPE aging.

## Methods

### Training and Testing Datasets

In this study, the mouse RPE flatmount images are selected from our reference database [19,20]. As these RPE images have high resolutions (around 4,000 × 4,000 pixels each), we divide each image into patches of size 96 × 96 in pixels. Both a small set of annotated and a large set of unlabeled RPE cells are included in our training set. For the small annotated set 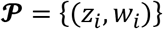, we have 155 patch pairs where *z_i_* and *w_i_* are the image patch and manually annotated ground truth. In the large set of unlabeled RPE cells for training, we include 987 image patches presenting RPE cells with weak or missing borders in 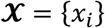, and 653 patches denoted by 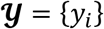 with strong cell borders. For the testing set, we include 34,742 RPE cells with weak or missing cell borders from 200 image patches.

### Deep Neural Network Architecture

For deep learning based image segmentation, deep neural networks often consist of an encoder and a decoder that are trained with a large amount of annotated data. By contrast, we make use of GAN-based image translation mechanism and use a semi-supervised learning strategy to improve the model performance due to a limited set of data annotations available in this work. In image translation tasks, GAN usually consists of two key components, i.e., a generator and a discriminator. The generator attempts to minimize the adversarial loss and translate inputs to images indistinguishable from real target images by the discriminator. By contrast, the discriminator is trained to maximize the adversarial loss and distinguish the fake images from real ones.

We present the overall architecture of the developed multi-head deep learning model (MultiHeadGAN) in Figure 2. Note MultiHeadGAN makes a full use of both limited data with annotations and a large set of unlabeled images for training. The generator in MultiHeadGAN is derived from the U-Net [18,21], but extended to multi-heads for contrast enhanced gray-scale and binary segmentation outputs. Different from the U-Net with one encoder and one decoder, our proposed generator has one encoder, two decoders, and one feature extractor. For each encoder input ***s***, there are two decoder output images *G*1(***s***) and *G*2(***s***) and a feature extractor output *V*(***s***). *G*1(***s***) from Decoder 1 represents a segmentation map, while *G*2(***s***) from Decoder 2 is the translated image with enhanced RPE cell borders. The output from the feature extractor *V*(***s***) is used for the contrastive representation learning in the model training. Although our generator has four resolution levels, not all levels are shown in Figure 2 for conciseness. At each image resolution level, the encoder convolves the image with a double convolution layer and next scales down the convolution response by a max-pooling layer. In the decoding analysis, an image representation is up-sampled, interpolated by a bilinear interpolation layer, and convolved with a double convolution layer in turn at each image resolution level. Additionally, the encoder outputs at different image resolution levels are processed by Multi-Layer Perceptron (MLP) modules, with the outcomes concatenated for the image feature vector construction.

**Figure 2.**
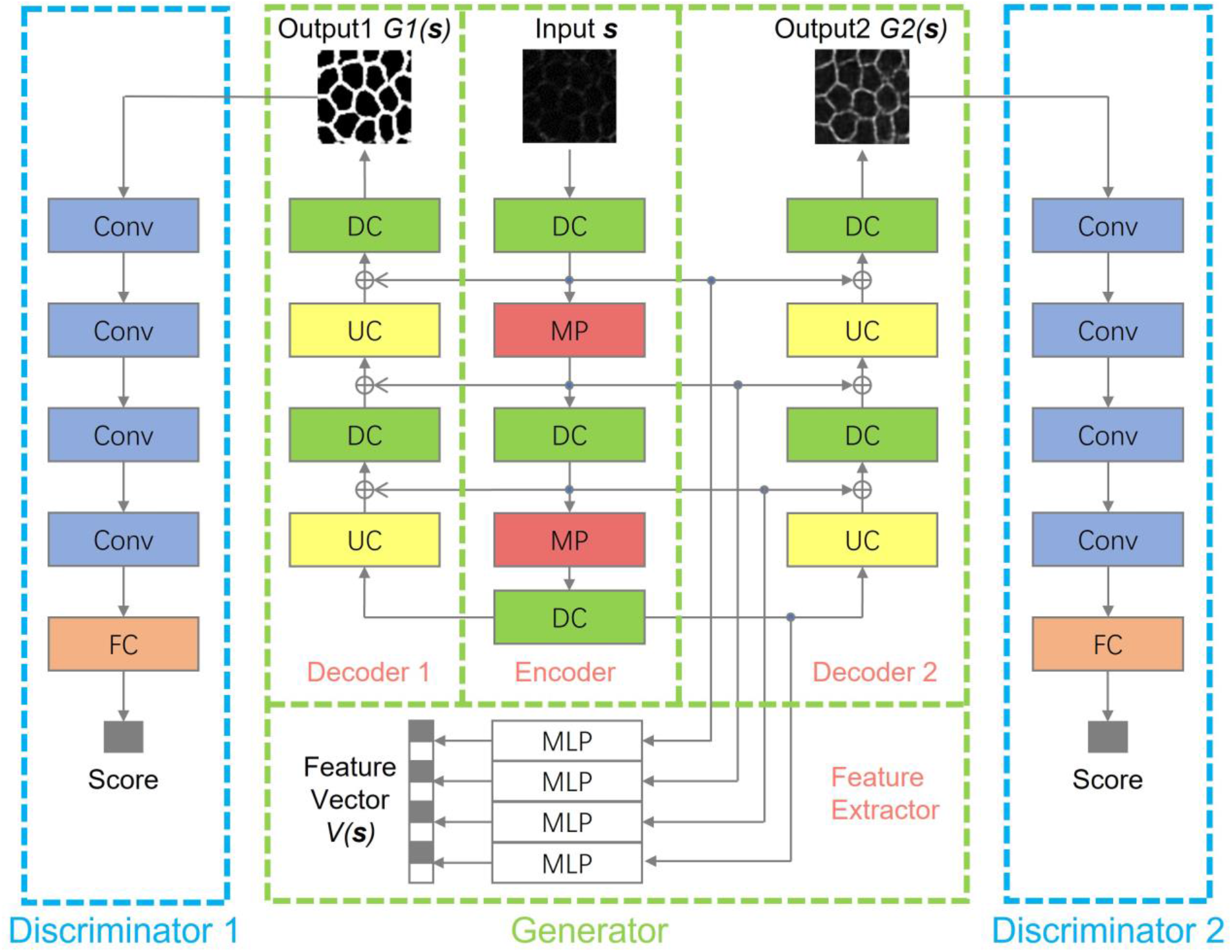
Overall schema of the developed multi-head deep learning approach MultiHeadGAN. Our proposed deep learning generator consists of one encoder, two decoders, and one feature extractor. For each image input, the network produces two output images and one feature vector. Note our generator has four image resolution levels. Not all levels are shown in the schema for conciseness. With such a model design, we can effectively detect RPE cell borders in damaged regions within flatmount microscopy images. *Conv*: Convolution layer; *DC*: Double convolution layers; *FC*: Fully connected layer; *MP*: Max-pooling layer; *MLP*: Multi-layer perceptron; *UC*: Up-sampling + convolution layer.

To process two image outputs *G*1(***s***) and *G*2(***s***) from the generator, we include two corresponding discriminators *D*1 and *D*2. By architecture, each discriminator has multiple convolution layers and a fully connected output layer [22]. These discriminators help recognize the difference between generated and true images and thus force the generator to produce high-quality images similar to the true counterparts.

### Model training strategy

With training batches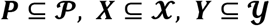, we would like to achieve two training objectives on image segmentation and translation. 1) For a given image and its ground truth pair (***z, w***)~***P***, we expect the segmentation result *G*1(***z***) from the generator to be similar to the segmentation ground truth ***w*** and indistinguishable by discriminator *D*1. 2) For RPE cells with weak (i.e. ***x***~***X***) and strong borders (i.e. ***y***~***Y***), we would like to make translated weak image *G*2(***x***) indistinguishable by discriminator *D*2 and keep translated strong images intact, i.e. *G*2(***y***) ≈ ***y***. To achieve these training goals, we define the objective function for the GAN training strategy as follows:

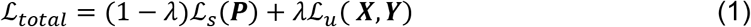

where two loss terms are balanced by the relative contribution factor *λ*.

This weight *λ* is dynamic and depends on the epoch number *t*:

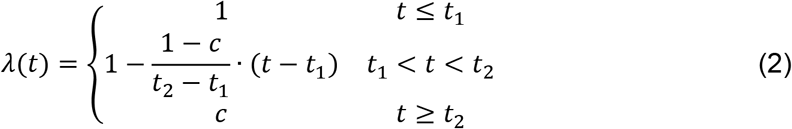

where *t*_1_ and *t*_2_ are transient time cutoff values; The constant *c* is the weight factor after *λ* is stabilized.

The first loss term 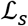 describes the similarity between the output of Decoder 1 and the segmentation ground truth:

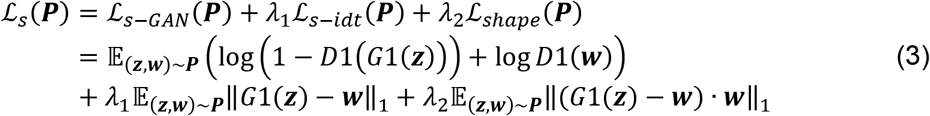

In Equation 3, the first two terms are the adversarial loss and the identity loss widely used in supervised GAN approaches [21,22]. In this work, our exploratory experimental results suggest that the segmented RPE cell borders are often not closed, leading to a significantly different RPE cell topology. This artifact results from the fact that the misclassification of cell border pixels has a small influence on the identity loss that in turn is due to a small proportion of cell border pixels in an entire image. In favor of closed RPE cell contours in the segmentation results, we penalize cell border misclassification more by a shape loss (i.e., the third term in Equation 3). In our designed shape loss term, we only direct model’s training attention to cell border misclassification events by multiplying the ground truth w to the difference between *G*1(***z***) and ***w***. As border and background pixels in the ground truth take value 1 and 0, respectively, such multiplication results in a focused attention to the misclassification on cell borders.

Similarly, the second loss term 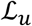 characterizes the quality of gray-scale outputs from the generator:

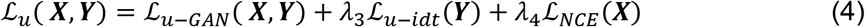

In Equation 4, the first term 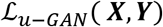 is the adversarial loss for unsupervised GAN learning that takes the following format:

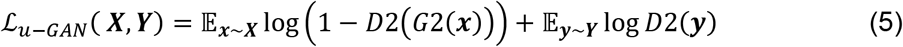

The second term 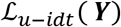 in Equation 4 is the identity loss to retain strong images at the generator output in the unsupervised image translation. While we aim to transfer weak to strong images by the generator, we also would like to keep those strong images unchanged during the translation. Therefore, the identity loss 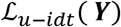 is defined by:

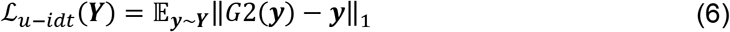

The third term 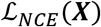 in Equation 4 is Noised Contrastive Estimation (NCE) loss [23]. It aims to train an encoder that associates only corresponding inputs [24]. In the unsupervised GAN training, this NCE loss prevents a generator from randomly producing images with high quality in the target domain but irrelevant to inputs [25]. Applying this loss to our work, we aim to achieve a high mutual information between an input ***x**_i_* and its translation output *G*2(***x**_i_*), and a low mutual information between the input ***x**_i_* and other translation outputs *G*2(***x**_j_*) Illustrated in Figure 2, encoded feature maps are processed with MLP modules for a vector representation *V*(***s***). Let ***v**_i_* = *V*(***x**_i_*) and 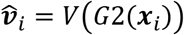, the NCE loss contributed by image ***x**_i_* is defined with a cross-entropy loss:

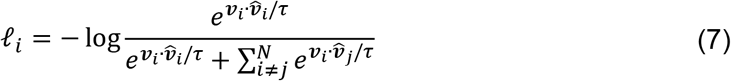

where *τ* is the scaling factor.

The resulting NCE loss for a training batch ***X*** is defined as:

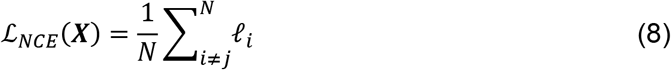

We implement our segmentation model with Python 3.8 programming language and PyTorch 1.8.1 deep learning framework [26] and run the segmentation analysis with two NVIDIA Tesla K80 GPUs. Balancing the tradeoff between computational efficiency and deep network efficacy, we design four image resolution levels in the generator, with 64, 128, 256, and 512 filters from the highest to the lowest level, respectively. Each MLP embedding image features from the Encoder has two neural network layers, with 256 units at each layer. For discriminators, we use two convolutional neural networks, both including the same resolution levels and filter numbers as the generator. Instead of max-pooling layers, image representations in discriminators are down-sampled by convolution layers of stride 2.

For training parameters, we have *t*_1_ = 40, *t*_2_ = 70, *c* = 0.7, *λ*_1_ = *λ*_2_ = 0.5, *λ*_3_ = *λ*_4_ = 1, and the scaling factor *τ* = 0.07 determined empirically.

### Evaluation metrics

In this work, we set border and background pixels as positive and negative classes, respectively. We derived metrics from the confusion matrix for model evaluations [16–18]. The confusion matrix results in TP (number of correctly classified border pixels), FP (number of incorrectly classified background pixels), FN (number of incorrectly classified border pixels), and TN (number of correctly classified background pixels). As TN is much larger than the other three in practice, we do not use accuracy ACC=(TP+TN)/(TP+FP+FN+TN)) as a metric. Instead, we adopt precision P=TP/(TP+FP) and recall R=TP/(TP+FN) for model evaluation.

Additionally, we introduce metrics to indicate RPE cell topology. For each cell with ground truth, we compute its Intersection over Union (IoU) with all overlapped cells predicted from models. If a cell has an IoU larger than 0.5, we mark it as a correct hit (CH); otherwise, it is a wrong hit (WH) as presented in Figure 3. We define the Correct Rate (CR) of segmentation as:

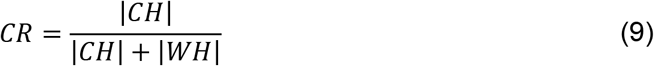

where |·| represents the set size.

**Figure 3.**
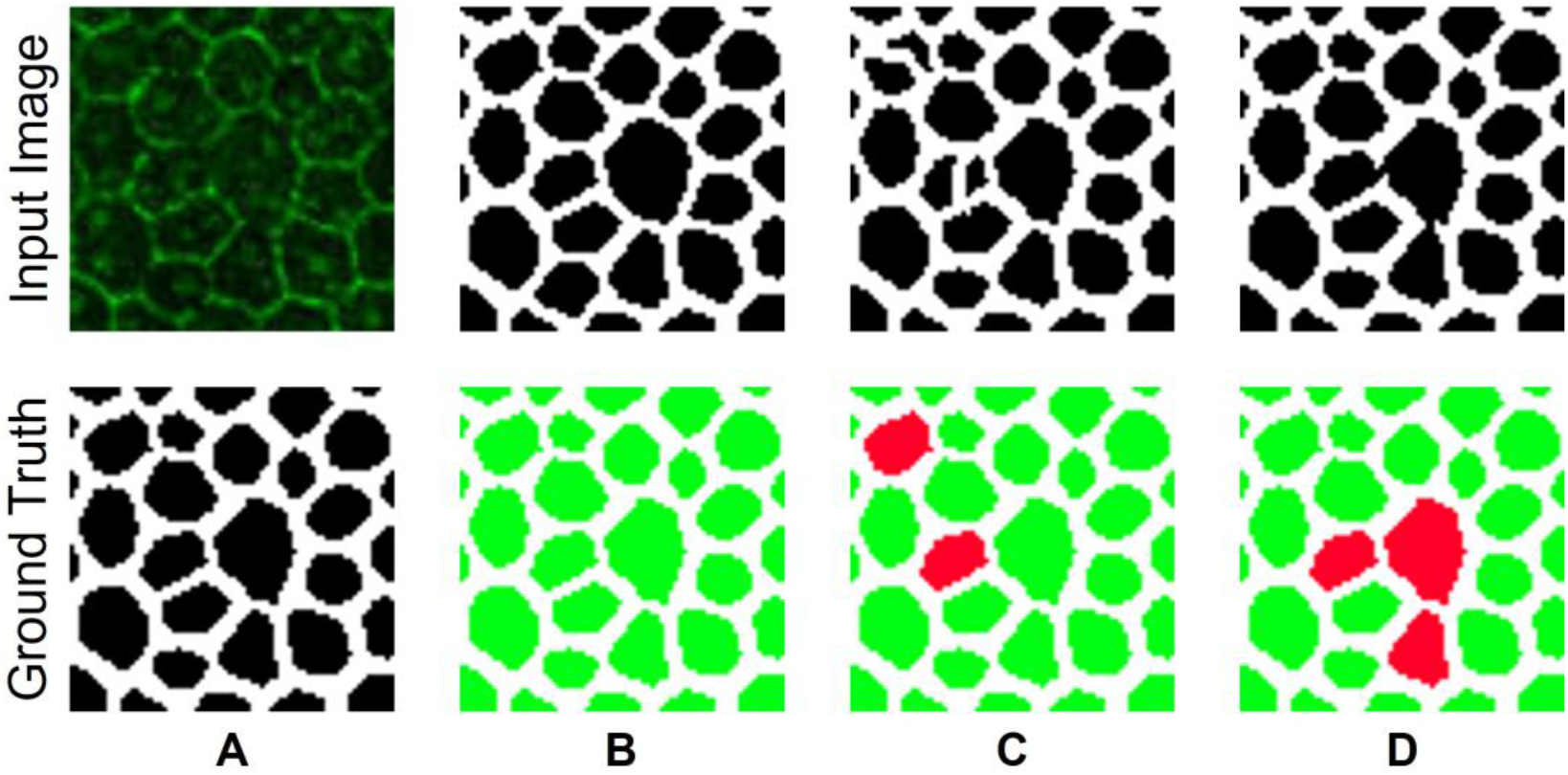
Evaluation of segmentation results. (A) A representative image (top) and its ground truth for segmentation (bottom) are presented. The (B) correct, (C) over-, and (D) under-segmentation results are illustrated, respectively. In (B-D), the top images present the segmentation results, while the bottom images highlight regions with erroneous segmentation results (red).

To penalize wrong segmentation by the cell size, we also use such information to calculate the Weighted Correct Rate (WCR) of segmentation:

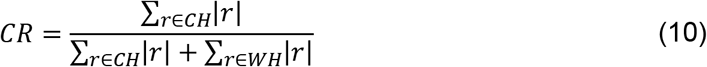

## Results

### Deep learning model validation

To validate our method performance, we compare our proposed method MultiHeadGAN with four state-of-the-art models, including FCN [17], DeepLab [16], UNet [18], and Cellpose [15]. FCN, DeepLab, and UNet have been widely applied to a large number of biomedical image segmentation applications [8]. Cellpose is a pre-trained cell segmentation model built on UNet. It is trained to predict gradient vector fields and produce segmentation results by gradient tracking. For fair comparisons, we use CycleGAN [22] and CUT [25] to enhance the RPE cell border contrast before UNet is used for segmentation. We train FCN, DeepLab and UNet with the annotated training set 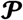 and trained CycleGAN and CUT with the unlabeled training sets 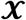 and 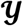. We present typical tissue segmentation results of these models in Figure 4. By visual comparisons, the results from FCN, DeepLab and UNet have large under-segmented regions. CycleGAN and CUT can effectively mitigate the degree of under-segmentation, while our developed MultiHeadGAN model achieves the best segmentation results. Although Cellpose can generate separated cell masks, its performance highly depends on the gradient vectors in cells. As not all cells present convergent gradient fields, Cellpose can fail in these cases.

**Figure 4.**
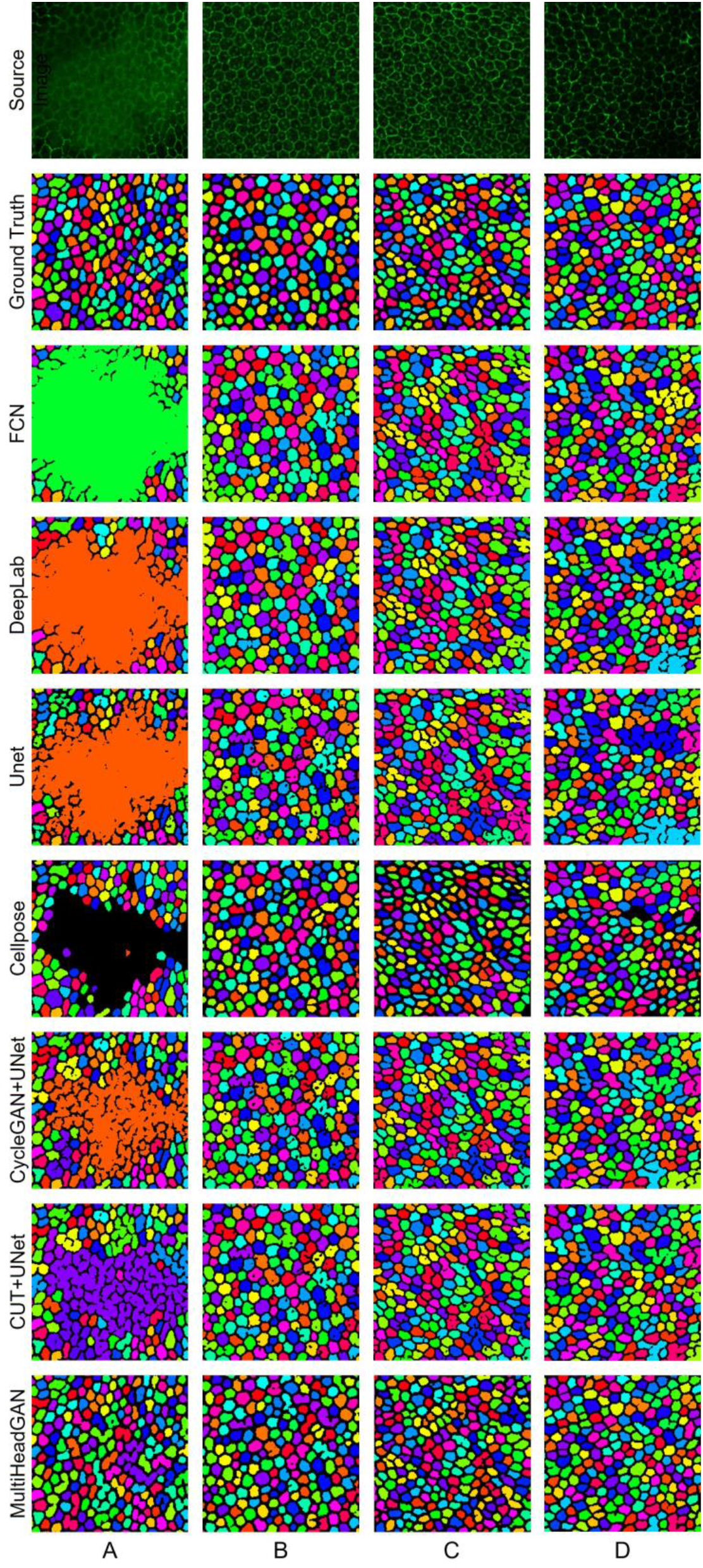
Qualitative comparison of deep learning approaches for RPE cell segmentation with flatmount microscopy images. Four typical impaired image regions are shown in columns with rows for ground truth and corresponding segmentation results of FCN, DeepLab, UNet, Cellpose, CycleGAN+UNet, CUT+UNet, and our developed MultiHeadGAN, respectively.

We quantitatively evaluate segmentation results by Correct Rate (CR), Weighted Correct Rate (WCR), Precision (P) and Recall (R). Demonstrated in Table 1, the proposed MultiHeadGAN achieves the best performance with 85.4% (CR), 88.8% (WCR), 87.3% (Precision) and 80.1% (Recall), respectively. In Figure 5, quantitative evaluation results are plotted to present the statistical difference between our proposed approach and others. Noticeably, MultiHeadGAN presents fewer outliers and a smaller variation, implying its strong stability. By metrics of WCR, P and R, MultiHeadGAN is significantly better than all other approaches. By CR, it is also significantly better than all other approaches except Cellpose.

**Table 1.** Comparison of quantitative performance of the developed MultiHeadGAN and other state-of-the-art deep learning approaches by different evaluation metrics on the mouse and human dataset. *L*: Training data with ground truth (i.e., 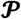); *UL*: Unlabeled training data (i.e., 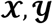); *CR*: Correct Rate; *WCR*: Weighted Correct Rate; *P*: Precision; *R*: Recall. All value units are in percentage (%).Included and excluded training data are checked by “✓” and “✗”, respectively. The absence is represented by “/”.

| Method |  | Training data |  | Mouse |  |  |  | Human |  |  |  |
| --- | --- | --- | --- | --- | --- | --- | --- | --- | --- | --- | --- |
| Enhancement | Segmentation | L | UL | CR | WCR | P | R | CR | WCR | P | R |
| / | FCN | ✓ | ✗ | 52.5 | 57.2 | 82.0 | 59.8 | 95.3 | 89.4 | 96.1 | **95.0** |
| / | DeepLab | ✓ | ✗ | 58.1 | 62.9 | 81.5 | 61.3 | 94.9 | **89.6** | 96.3 | 94.9 |
| / | UNet | ✓ | ✗ | 56.1 | 60.1 | 85.0 | 68.5 | 92.4 | 85.2 | 97.9 | 90.6 |
| / | Cellpose | / | / | 83.8 | 86.7 | 73.8 | 73.7 | **97.1** | 87.9 | 95.1 | 91.4 |
| CycleGAN | UNet | ✓ | ✓ | 59.9 | 64.2 | 85.7 | 70.0 | 96.7 | 88.8 | **98.9** | 92.4 |
| CUT | UNet | ✓ | ✓ | 64.5 | 68.8 | 86.1 | 70.3 | 96.9 | 89.0 | **98.9** | 93.0 |
| MultiHeadGAN |  | ✓ | ✓ | **85.4** | **88.8** | **87.3** | **80.1** | 96.5 | 88.8 | **98.9** | 92.7 |

**Figure 5.**
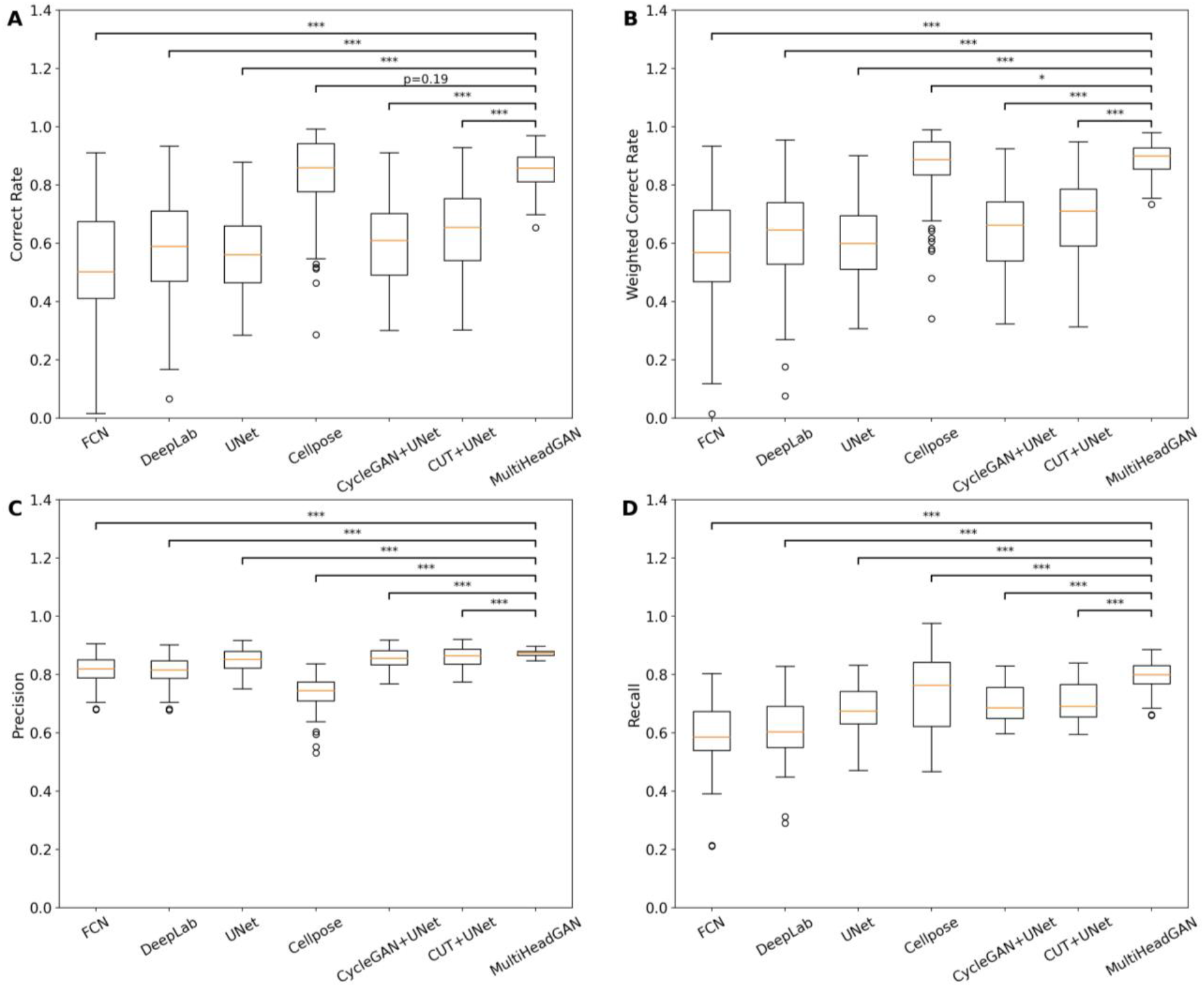
Quantitative comparison of deep learning approaches for RPE flatmount image segmentation. We compare the RPE cell segmentation performance of deep learning approaches by (A) Correct Rate, (B) Weighted Correct Rate, (C) Precision, and (D) Recall. Paired sample t-tests between the developed MultiHeadGAN and other six state-of-the-art approaches suggest a statistically significant performance difference. The notations for *, **, and *** represent a p-value less than 0.05, 0.005, and 0.0005, respectively.

**Figure 6.**
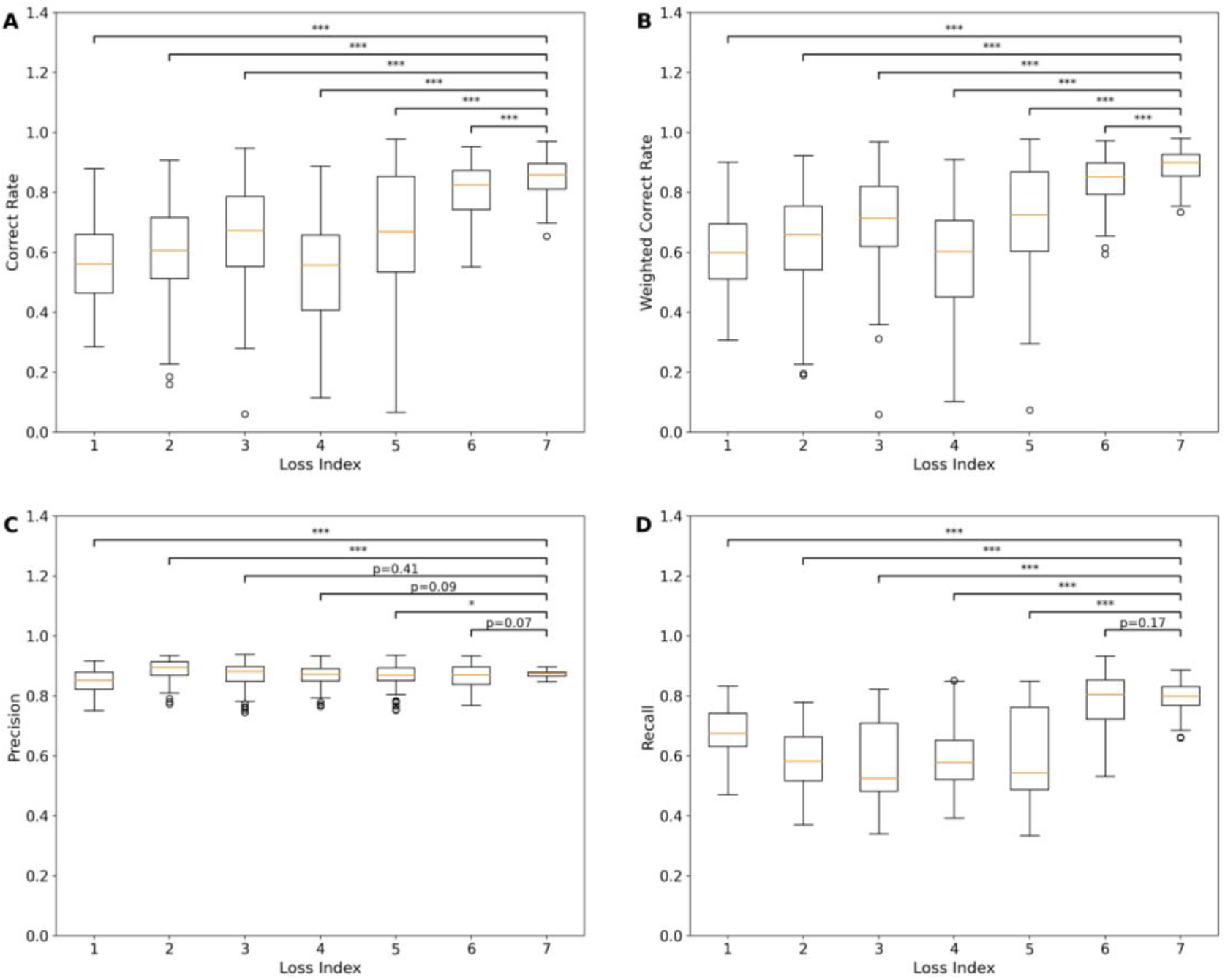
Quantitative comparison of ablated models for RPE flatmount image segmentation. We compare the segmentation performance of our model with different training loss combinations by (A) Correct Rate, (B) Weighted Correct Rate, (C) Precision, and (D) Recall. The x-axis represents the loss combination index given in Table 2. Paired sample t-tests between the proposed combination for MultiHeadGAN and other six loss combinations suggest a statistically significant performance difference in most cases. The notations for *, **, and *** represent a p-value less than 0.05, 0.005, and 0.0005, respectively.

Additionally, we test all approaches on a human RPE flatmount image dataset. Each image has 1,024×1,024 pixels by size. The training and testing data include 14 and 16 human samples. We use the human training set for transfer learning with FCN, DeepLab, UNet and MultiHeadGAN. The resulting method performances are shown in Table 1. As most cell borders in our human dataset are strong, all methods for comparison present no significant difference. Our proposed approach achieves the best performance by Precision (98.9%). By CR, WCR, and Recall, all methods present similar performances. The difference between our proposed approach and the best approach is only 0.6%, 0.6%, and 2.3%, respectively.

### Ablation study

To demonstrate the contribution from individual training losses in our developed MultiHeadGAN model, we carry out ablation experiments with seven loss combinations. The resulting performances are presented in Table 2. Loss combination 7 is for our MultiHeadGAN model with all losses and achieves the best performance. Comparisons between loss combination 1 and 2 imply that the GAN mechanism improves supervised training outcome. Furthermore, comparisons between loss combination 2 and 3 suggest that our proposed shape loss can boost the performance with annotated training data. Comparisons between loss combination 2 and 4 indicate that the addition of unsupervised learning to deep learning model training may degrade the model performance without use of appropriate constraints (i.e., identity loss and NCE loss).

**Table 2.** Quantitative performance comparison across different training loss combinations by different evaluation metrics on the mouse dataset. *CR*: Correct Rate; *WCR*: Weighted Correct Rate; *P*: Precision; *R*: Recall. All value units are in percentage (%). Included and excluded loss terms are checked by “✓” and “✗”, respectively.

| Loss Index | Loss Term |  |  |  |  |  | CR | WCR | P | R |
| --- | --- | --- | --- | --- | --- | --- | --- | --- | --- | --- |
| | $\mathcal{L}_{S-idt}$ | $\mathcal{L}_{S-GAN}$ | $\mathcal{L}_{u-GAN}$ | $\mathcal{L}_{u-idt}$ | $\mathcal{L}_{NCE}$ | $\mathcal{L}_{shape}$ | | | | |
| 1 | ✓ | ✗ | ✗ | ✗ | ✗ | ✗ | 56.1 | 60.1 | 85.0 | 68.5 |
| 2 | ✓ | ✓ | ✗ | ✗ | ✗ | ✗ | 59.4 | 63.4 | **88.7** | 58.7 |
| 3 | ✓ | ✓ | ✗ | ✗ | ✗ | ✓ | 66.4 | 70.5 | 87.0 | 57.4 |
| 4 | ✓ | ✓ | ✓ | ✗ | ✗ | ✗ | 53.4 | 57.2 | 86.8 | 60.4 |
| 5 | ✓ | ✓ | ✓ | ✓ | ✗ | ✗ | 67.7 | 71.3 | 86.7 | 59.4 |
| 6 | ✓ | ✓ | ✓ | ✓ | ✓ | ✗ | 80.9 | 84.2 | 86.7 | 78.9 |
| 7 | ✓ | ✓ | ✓ | ✓ | ✓ | ✓ | **85.4** | **88.8** | 87.3 | **80.1** |

Note we have a dynamic weight factor *λ* to balance the supervised and unsupervised learning. To validate the effectiveness of this design and determine corresponding parameters, we have carried out ablation experiments. In Figure 7, the training curves of identity loss for the supervised learning suggest that the parameter setting with *t*_1_ = 40, *t*_2_ = 70, *c* = 0.7 achieves the best training performance. Furthermore, we compare the testing performance in Table 3.

**Figure 7.**
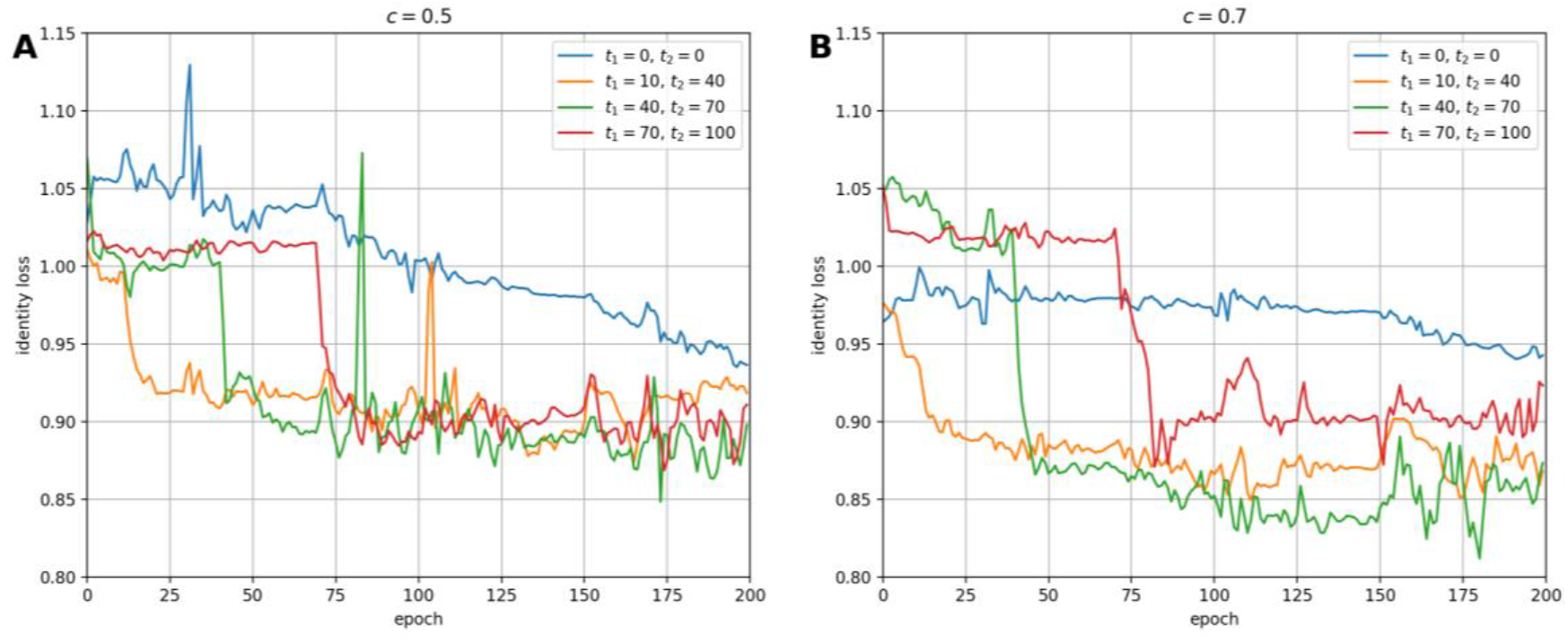
Comparison of our model training curves with different training parameters. We compare the training curves of the identity loss for the supervised learning with different strategies for the weight factor *λ*. Training curves with stabilized weight factor *c* = 0.5 and *c* = 0.7 are presented in (A) and (B), respectively.

**Table 3.**
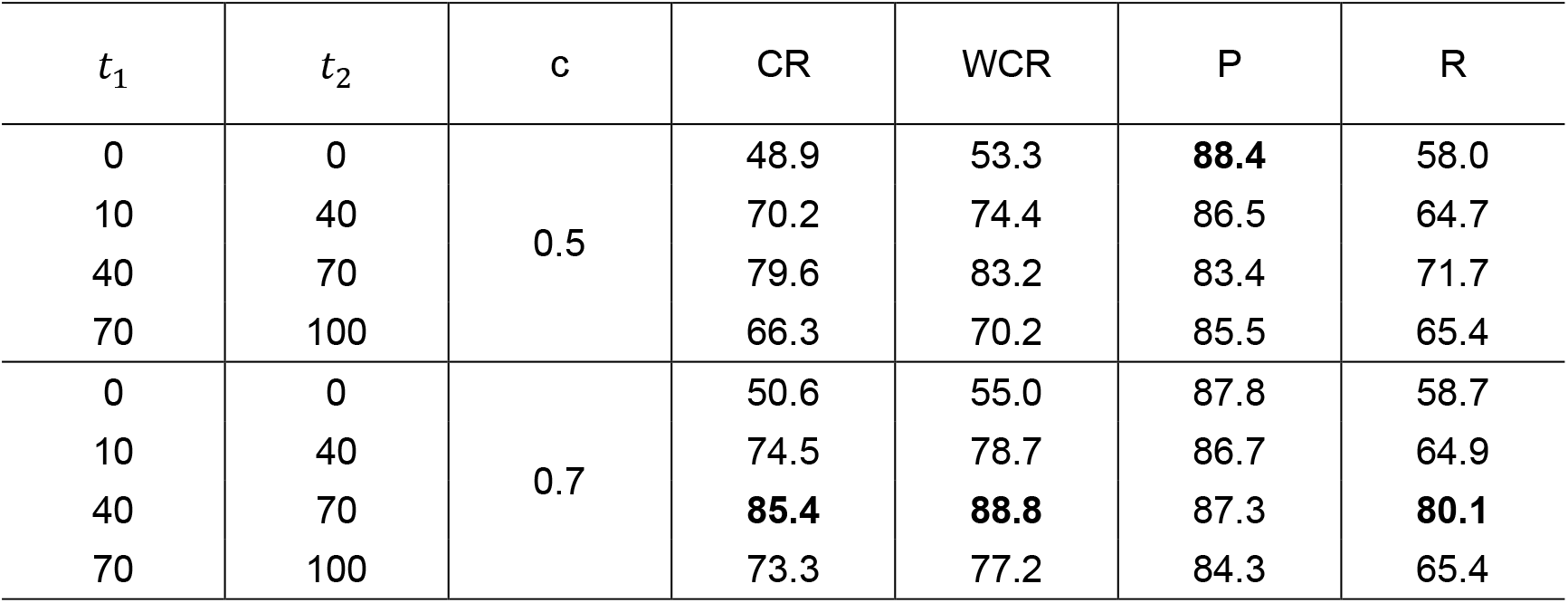
Quantitative performance comparison with different weight factor *λ* strategies by different evaluation metrics on the mouse dataset. *CR*: Correct Rate; *WCR*: Weighted Correct Rate; *P*: Precision; *R*: Recall. All value units are in percentage (%).

**Table 4.** Comparison of deep learning segmentation models by model parameter number, Floating Point Operations Per Second (FLOPS), and the average processing time cost on a 96×96 image patch.

| Model | Parameter number | FLOPS | Processing time (ms)* |
| --- | --- | --- | --- |
| FCN | 51.9M | 7.6G | 134.6 |
| DeepLab | 58.6M | 8.5G | 167.8 |
| UNet | 34.5M | 9.2G | 92.9 |
| Cellpose | 6.6M | 2.9G | 3,603.6 |
| CycleGAN+UNet | 43.0M | 16.1G | 132.6 |
| CUT+UNet | 43.0M | 16.0G | 125.1 |
| MultiHeadGAN | 8.5M | 7.0G | 35.9 |

With a constant weight factor *λ* (i.e., *t*_1_ = *t*_2_ = 0), our model is trained with both supervised and unsupervised learning. The results suggest that a fixed *λ* has the worst performance across dynamic factor strategies for each given stabilized weight factor *c*. Next, we begin model training only with the unsupervised learning and linearly change the weight factor *λ* for a semi-supervised learning. We define the pre-training as the stage only with unsupervised learning. The performance is improved as the pre-training time increases. When the pre-training time is increased to 40 epochs, our model achieves the best performance. When the pre-training time is increased further, our model performance becomes worse in large part due to overfitting in the unsupervised learning. After systematic investigations, we choose parameters *t*_1_ = 40, *t*_2_ = 70, *c* = 0.7 for the dynamic weight factor strategy by the experiment results.

## Discussion

RPE cell morphological information plays a vital role to facilitate a better understanding of RPE aging and physiology. An accurate morphological characterization, however, is highly dependent on a high-quality segmentation. Due to lack of appropriate computational methods, the segmentation maps are often created by manual annotations, suffering from a large inter- and intra-variability. Additionally, such a human annotation process is time-consuming and insufficient to produce an adequately large number of annotated RPE images from impaired regions to support computer-based method training.

To address this challenge, we develop a novel method (MultiHeadGAN) for segmenting RPE cells from 2D RPE flatmount microscopy images in this study. Our developed method takes a semi-supervised learning strategy and enables deep neural networks to learn from a small set of annotated and a large set of unlabeled image data, resulting in a more generalized feature extraction ability and learning outcome. The resulting deep neural network is designed within a GAN training framework equipped with one encoder, two decoders, and one feature extractor in the generator. Two heads are created in the generator for the contrast enhanced grayscale and binary segmentation output, respectively. Correspondingly, there are two discriminators in our model that force the generator to output images with strong borders. Additionally, we create a new shape loss term that encourages our model to produce closed cell borders.

Demonstrated in Figure 3, very few mis-classified pixels can either connect separate cells or divide a single cell, leading to a huge change in the RPE cell topology. We propose two new metrics, i.e. Correct Rate (CR) and Weighted Correct Rate (WCR), to better characterize such cell topology. CR represents the proportion of correctly segmented cells in the total cell population, while WCR indicates the proportion of areas of correctly segmented cells in the total cell area. As shown in Figure 5, the differences across different methods by CR and WCR are much larger than those suggested by Precision and Recall. CR and WCR can better reflect performance difference supported by the visual results in Figure 4.

We have systematically compared our MultiHeadGAN model with other state-of-the-art deep learning methods. MultiHeadGAN manifests a superior performance by RPE cell segmentation accuracy, model complexity, and computational efficiency. By both visual and quantitative evaluations, we demonstrate the promising segmentation performance by MultiHeadGAN. In the ablation study, we systematically compare and analyze the contribution from each loss term. Although our approach takes a semi-supervised training strategy for multiple network components in a GAN framework, we only need to load Encoder and Decoder 1 in the testing stage. MultiHeadGAN includes the least parameters and requires the least Floating Point Operations Per Second (FLOPS) for processing except Cellpose. Although Cellpose has the least number of parameters, it requires a time-consuming gradient tracking process, resulting in the longest average processing time (3,604 ms). Additionally, models of CycleGAN+Unet and CUT+Unet leverage the GAN mechanism and include more parameters, leading to a slower analysis. By contrast, our proposed MultiHeadGAN has the shortest average processing time cost (35.9 ms), promising to support large-scale RPE image data analysis.

In future work, we plan to aggregate “network engineering” techniques in the generator to boost the segmentation performance further. Residual connections have demonstrated its potential for biomedical image segmentation [13,27]. Inclusion of such residual connections to our network may improve image feature information learning. Additionally, attention layers are known to encourage networks to focus on features in target regions [28]. Thus, addition of attention layers to our generator may better guide it for RPE cell border feature extraction. Furthermore, depth-wise separable convolution layers can be used to replace regular convolution layers, as they can help reduce model complexity and accelerate training speed [29,30]. As Transformer based networks have achieved state-of-the-art results in some medical image segmentation tasks [7], we also plan to make them as new generators in our project. Finally, we plan to generalize our approach to support a larger set of biomedical image based disease investigations where cell borders are weak but critical for morphology analyses.

## Conclusion

A new approach MultiHeadGAN has been developed to segment RPE cells with weak or missing borders in damaged regions of flatmount microscopy images, allowing more informative and reliable cell morphology investigations. Built upon the UNet architecture, we use one encoder and two separate decoders to enable simultaneous training with both limited data with human annotations and a large set of unlabeled images by the GAN mechanism. The performance of the model is systematically validated and compared with other state-of-the-art deep learning approaches both in a qualitative and quantitative manner. Our developed model makes it possible to segment low contrast RPE cells in flatmount microscopy images without expensive annotated training datasets, presenting its promising potential to assist cell morphology analyses for RPE aging studies.

## Data Availability Statement

Codes are available at Github repository: https://github.com/jkonglab/RPE_MultiHeadGAN Testing images are available at https://figshare.com/s/0040cad8211ed9517072

## Acknowledgments

This research is supported in part by grants from National Institute of Health [1U01CA242936, R01EY028450, R01EY021592, P30EY006360, R01EY028859, T32EY07092, T32GM008490]; National Science Foundation [ACI 1443054 and IIS 1350885]; the Abraham J. and Phyllis Katz Foundation; the U.S. Department of Veterans Affairs and Atlanta Veterans Administration Center for Excellence in Vision and Neurocognitive Rehabilitation [RR&D I01RX002806, I21RX001924; VA RR&D C9246C]; CNPq and FAPEMIG agencies.

## References

[1] Strauss, O. “The retinal pigment epithelium in visual function.” Physiological reviews, vol. 85, no. 3, 2005, pp. 845–81.

[2] Ambati, J., Ambati, B.K., Yoo, S.H. et al. “Age-related macular degeneration: etiology, pathogenesis, and therapeutic strategies.” Survey of ophthalmology, vol. 48, no. 3, 2003, pp. 257–93.

[3] Ach, T., Huisingh, C., McGwin, G. et al. “Quantitative autofluorescence and cell density maps of the human retinal pigment epithelium.” Investigative ophthalmology & visual science, vol. 55, no. 8, 2014, pp. 4832–41.

[4] Bhatia, S.K., Rashid, A., Chrenek, M.A. et al. “Analysis of RPE morphometry in human eyes.” Molecular vision, vol. 22, 2016, p. 898.

[5] Kim, Y.K., Yu, H., Summers, V.R. et al. “Morphometric Analysis of Retinal Pigment Epithelial Cells From C57BL/6J Mice During Aging.” Invest Ophthalmol Vis Sci, vol. 62, no. 2, 2021, p. 32, doi:10.1167/iovs.62.2.32.

[6] Arganda-Carreras, I., Kaynig, V., Rueden, C. et al. “Trainable Weka Segmentation: a machine learning tool for microscopy pixel classification.” Bioinformatics, vol. 33, no. 15, 2017, pp. 2424–26.

[7] Shamshad, F., Khan, S., Zamir, S.W. et al. “Transformers in Medical Imaging: A Survey.” arXiv preprint arXiv:2201.09873, 2022.

[8] Zhou, S.K., Greenspan, H., Davatzikos, C. et al. “A review of deep learning in medical imaging: Imaging traits, technology trends, case studies with progress highlights, and future promises.” Proceedings of the IEEE, vol. 109, 2021, pp. 820–38.

[9] Guo, X., Wang, F., Teodoro, G. et al. “Liver steatosis segmentation with deep learning methods.” 2019 IEEE 16th International Symposium on Biomedical Imaging, IEEE, 2019, pp. 24–27.

[10] Zeng, Z., Xie, W., Zhang, Y. et al. “RIC-Unet: An improved neural network based on Unet for nuclei segmentation in histology images.” Ieee Access, vol. 7, 2019, pp. 21420–28.

[11] Li, X., Chen, H., Qi, X. et al. “H-DenseUNet: hybrid densely connected UNet for liver and tumor segmentation from CT volumes.” IEEE transactions on medical imaging, vol. 37, no. 12, 2018, pp. 2663–74.

[12] Wang, C., MacGillivray, T., Macnaught, G. et al. “A two-stage 3D Unet framework for multi-class segmentation on full resolution image.” arXiv preprint arXiv:1804.04341, April 12, 2018 2018. https://arxiv.org/abs/1804.04341.

[13] Ibtehaz, N. and Rahman, M.S. “MultiResUNet: Rethinking the U-Net architecture for multimodal biomedical image segmentation.” Neural Networks, vol. 121, 2020, pp. 74–87.

[14] Greenwald, N.F., Miller, G., Moen, E. et al. “Whole-cell segmentation of tissue images with human-level performance using large-scale data annotation and deep learning.” bioRxiv, 2021.

[15] Stringer, C., Wang, T., Michaelos, M. et al. “Cellpose: a generalist algorithm for cellular segmentation.” Nature Methods, vol. 18, no. 1, 2021, pp. 100–06.

[16] Chen, L.-C., Papandreou, G., Kokkinos, I. et al. “Deeplab: Semantic image segmentation with deep convolutional nets, atrous convolution, and fully connected CRFs.” IEEE transactions on pattern analysis and machine intelligence, vol. 40, no. 4, 2017, pp. 834–48.

[17] Long, J., Shelhamer, E., and Darrell, T. “Fully convolutional networks for semantic segmentation.” Proceedings of the IEEE conference on computer vision and pattern recognition, IEEE, 2015, pp. 3431–40.

[18] Ronneberger, O., Fischer, P., and Brox, T. “U-Net: Convolutional networks for biomedical image segmentation.” International Conference on Medical image computing and computer-assisted intervention, Springer, 2015, pp. 234–41.

[19] Boatright, J.H., Dalal, N., Chrenek, M.A. et al. “Methodologies for analysis of patterning in the mouse RPE sheet.” Molecular vision, vol. 21, 2015, p. 40.

[20] Jiang, Y., Qi, X., Chrenek, M.A. et al. “Functional principal component analysis reveals discriminating categories of retinal pigment epithelial morphology in mice.” Investigative ophthalmology & visual science, vol. 54, no. 12, 2013, pp. 7274–83.

[21] Isola, P., Zhu, J.-Y., Zhou, T. et al. “Image-to-image translation with conditional adversarial networks.” Proceedings of the IEEE conference on computer vision and pattern recognition, 2017, pp. 1125–34.

[22] Zhu, J.-Y., Park, T., Isola, P. et al. “Unpaired image-to-image translation using cycle-consistent adversarial networks.” Proceedings of the IEEE international conference on computer vision, 2017, pp. 2223–32.

[23] Gutmann, M. and Hyvärinen, A. “Noise-contrastive estimation: A new estimation principle for unnormalized statistical models.” Proceedings of the thirteenth international conference on artificial intelligence and statistics, JMLR Workshop and Conference Proceedings, 2010, pp. 297–304.

[24] Oord, A.v.d., Li, Y., and Vinyals, O. “Representation learning with contrastive predictive coding.” arXiv preprint arXiv:1807.03748, 2018.

[25] Park, T., Efros, A.A., Zhang, R. et al. “Contrastive learning for unpaired image-to-image translation.” European Conference on Computer Vision, Springer, 2020 2020, pp. 319–45.

[26] Paszke, A., Gross, S., Massa, F. et al. “Pytorch: An imperative style, high-performance deep learning library.” Advances in neural information processing systems, vol. 32, 2019, pp. 8026–37.

[27] Drozdzal, M., Vorontsov, E., Chartrand, G. et al. “The Importance of Skip Connections in Biomedical Image Segmentation.” Deep Learning and Data Labeling for Medical Applications, edited by Gustavo Carneiro et al., Springer International Publishing, 2016, pp. 179–87.

[28] Woo, S., Park, J., Lee, J.-Y. et al. “CBAM: Convolutional block attention module.” Proceedings of the European conference on computer vision (ECCV), Springer, 2018, pp. 3–19.

[29] Lee, J.H., Joo, I., Kang, T.W. et al. “Deep learning with ultrasonography: automated classification of liver fibrosis using a deep convolutional neural network.” European radiology, vol. 30, no. 2, 2020, pp. 1264–73.

[30] Howard, A.G., Zhu, M., Chen, B. et al. “MobileNets: Efficient convolutional neural networks for mobile vision applications.” arXiv preprint arXiv:1704.04861, April 17, 2017 2017. https://arxiv.org/abs/1704.04861.

